# Chromosome-level genome assembly of *Tamarindus indica*

**DOI:** 10.1101/2025.04.01.646601

**Authors:** Mitali Singh, Manohar S. Bisht, M G Abhijith, Shruti Mahajan, Vineet K. Sharma

**Affiliations:** MetaBioSys Group, Department of Biological Sciences, Indian Institute of Science Education and Research Bhopal, Bhopal, India

**Keywords:** Chromosome level assembly, Tamarind, Whole genome duplication, Leguminosae

## Abstract

*Tamarindus indica* is the sole member of the genus *Tamarindus* of the Leguminosae family. It is a multipurpose horticultural plant, with every part of the plant finding importance in food, medicine, and other industries. To gain an understanding of genome structure and evolution, we reported the first high-quality genome assembly of *T. indica* anchored to 12 chromosomes with N50 of ∼56 Mb. Supported by comprehensive transcriptome data, we reported >48,000 protein-coding genes. Through phylogenetic and evolutionary analysis, we uncovered an independent whole-genome duplication event in *T. indica* and highlighted the expression divergence of segmentally duplicated genes and their role in the better adaptivity of the plant. Our study thus provides first insights into its genomic organization, evolutionary history, and WGD event and thus becomes an important resource for future genetic and biotechnological studies to understand essential pathways and assist breeding programs for trait enhancements.

## Introduction

*Tamarindus indica,* commonly known as tamarind, is a long-lived eudicot plant belonging to the family Leguminosae and a sole member of the genus *Tamarindus* (1). Its name originated from the Arabic word “Tamar ul’ Hind,” meaning dates of India. It is believed to be native to Africa and distributed to other tropical Asian countries, especially India, thereby getting its name. It is widely distributed and cultivated across South Asian, African, and American countries (2). Almost all parts of the tamarind plant hold significance due to its medicinal, industrial, and culinary use (1,2). The plant is renowned for its high nutraceutical value, attributed to the prominent level of dietary fibers, vitamins (like folates, niacin, thiamine, Vitamin A, C, and E), electrolytes (sodium, potassium), and minerals (2,3). Additionally, it contains various organic acids (tartaric acid, malic acid, citric acid, etc.) and other phytonutrients (4,5). The phytochemicals found in the different tissues impart several health and medicinal benefits, including antidiabetic, antimicrobial, antioxidant, hepatoprotective, and anti-inflammatory properties (4).

Metabolic explorations of the extracts of this plant have shown significant contents of triterpenoids and phytosterols. Two important triterpenoids, Lupeol and Lupanone, were detected and isolated from the leaves of *T. indica* (6). Lupeol content was also found to be high in the flowers and bark of the plant, which imparts anti-microbial and antioxidant properties, contributing to the plant’s defense mechanism and overall health (7). Besides this, it has emerged as a potent dietary supplement with immense therapeutic value (8,9).

Despite having significant nutraceutical and pharmacological value, *T. indica* remains a highly underutilized plant, with its applications mainly confined to delicacies, flavoring agents, and traditional medicines. The absence of genomic resources limits the exploration and translational application of its essential biosynthetic pathways. In this study, we generated the first chromosome-level genome of *T. indica* (2n=24). We carried out an in-depth analysis of its phylogenomic and gene family evolution dynamics and examined the role of segmentally duplicated genes and their expression divergence. These findings will provide advances in both biological and evolutionary perspectives while also elucidating the genome structure and organization of *T. indica*.

## Results

### Genome assembly and quality assessment

A total of 78x coverage of linked reads data from 10x Genomics, 28x of Oxford Nanopore reads, ∼81 Gb of transcriptome data (>15Gb per tissue), and ∼200 million paired-end Hi-C reads were generated. Through K-mer analysis of the adapter-trimmed 10x Genomics linked reads, the genome size of *T. indica* was estimated to be 778 Mb, with a heterozygosity of 1.12%. The merged assemblies of the 10x Genomics linked reads, and Oxford Nanopore long reads resulted in an assembled genome of size 850.4 Mb, with a contig N50 of 4.0 Mb. Further genome scaffolding was carried out using genome-wide chromatin interaction information obtained by sequencing the Hi-C library. The contact map generated from the Hi-C reads resulted in anchoring the genome into 12 superscaffolds (chromosomes) corresponding to ∼93% of the total assembled genome (**Figure 1A and 1B**). The final genome assembly of *T. indica* was of size 776 Mb, which is near the estimated genome size with a scaffold N50 of ∼56 Mb and the largest scaffold of size 88 Mb.

**Figure 1:**
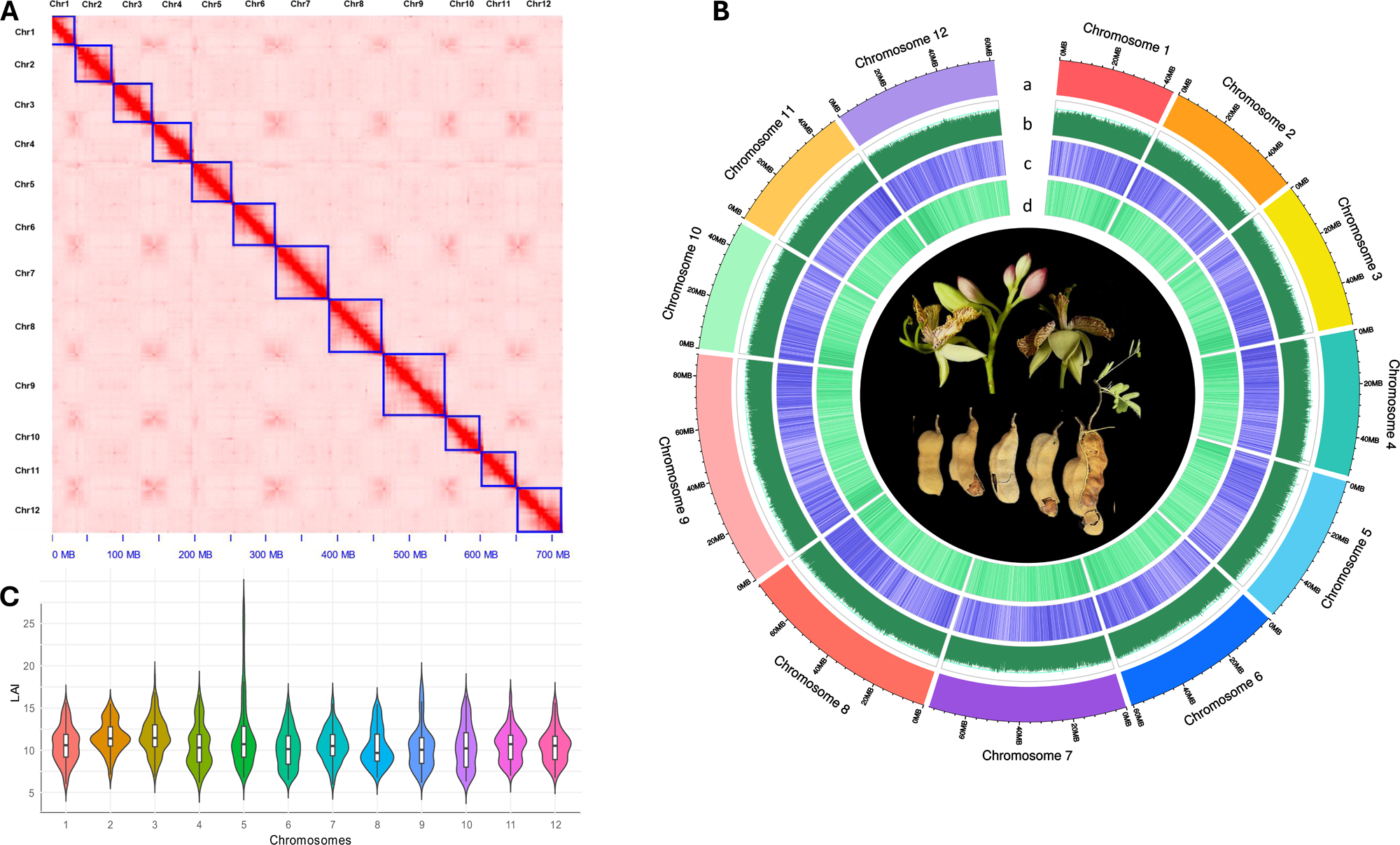
Characteristics of *T. indica* genome assembly. **A.** Chromosomal interaction heatmap: Blue frames indicate the 12 assembled chromosomes, **B.** Circos plot of *T. indica* genome: The tracks from outside to inside are (a) 12 chromosomes length, (b) GC content, (c) Repeat density, and (d) Gene density. All distributions are drawn in a window size of 100 kb, **C.** LAI scores distribution of 12 chromosomes of *T. indica*.

For the assessment of the quality and completeness of the genome assembly, the mapping percentage of the raw reads to the final assembly and the BUSCO score of the assembly were calculated. 95.58% of the 10x linked reads and 95.16% of Nanopore long reads could be aligned to the final genome assembly. The BUSCO completeness score for the final genome assembly was 92.8%. Also, the long terminal repeat (LTR) assembly index (LAI) score of all the 12 chromosomes had a value close to or above 10 (**Figure 1C**), with the final genome assembly LAI of 10.84, suggesting its quality to be of reference standard.

### Gene set construction and annotation

The *de novo* repeat library identified 63.85% of repetitive sequences in the genome. Among the interspersed repeats, Gypsy/DIRS1 constituted 25.22%, and Ty1/Copia constituted 10.51%. Combining homology and *ab initio* based gene prediction models, a total of 48,867 protein-coding genes were predicted, of which 93.7% could be functionally annotated using publicly available databases. Noncoding RNA prediction identified 418 miRNAs, 575 tRNAs, and 796 rRNAs. The mapping of the transcripts to the final genome assembly resulted in the alignment of 93.95%, 84.31%, 90.18%, and 89.62% from the leaf, flower buds, fruit pulp, and seed tissues, respectively.

### Phylogenetic tree construction and gene family evolution

A total of 531 fuzzy one-to-one orthgroups were identified from the selected 20 plant species and were used for phylogenetic analysis. The phylogenetic tree placed *T. indica* closest to *Cercis canadensis*. *Tamarindus* belongs to the Detarioideae subfamily, which diverged early along with the Cercidoideae subfamily from the other legumes. Later, it diverged from the Cercidoideae subfamily around the estimated divergence time of 58.12 mya (**Figure 2A)**. The species phylogenetic tree constructed in this study is consistent with the Legume Phylogeny Working Group (LPWG) (10). CAFÉ analysis for the gene family expansion and contraction resulted in the identification of 14,280 filtered gene families across the selected species. Of the 8,328 gene families of *T. indica*, 1,484 gene families were expanded, and 3,278 gene families were contracted. Among the expanded gene families, 12 gene families were highly expanded (>10 genes).

**Figure 2:**
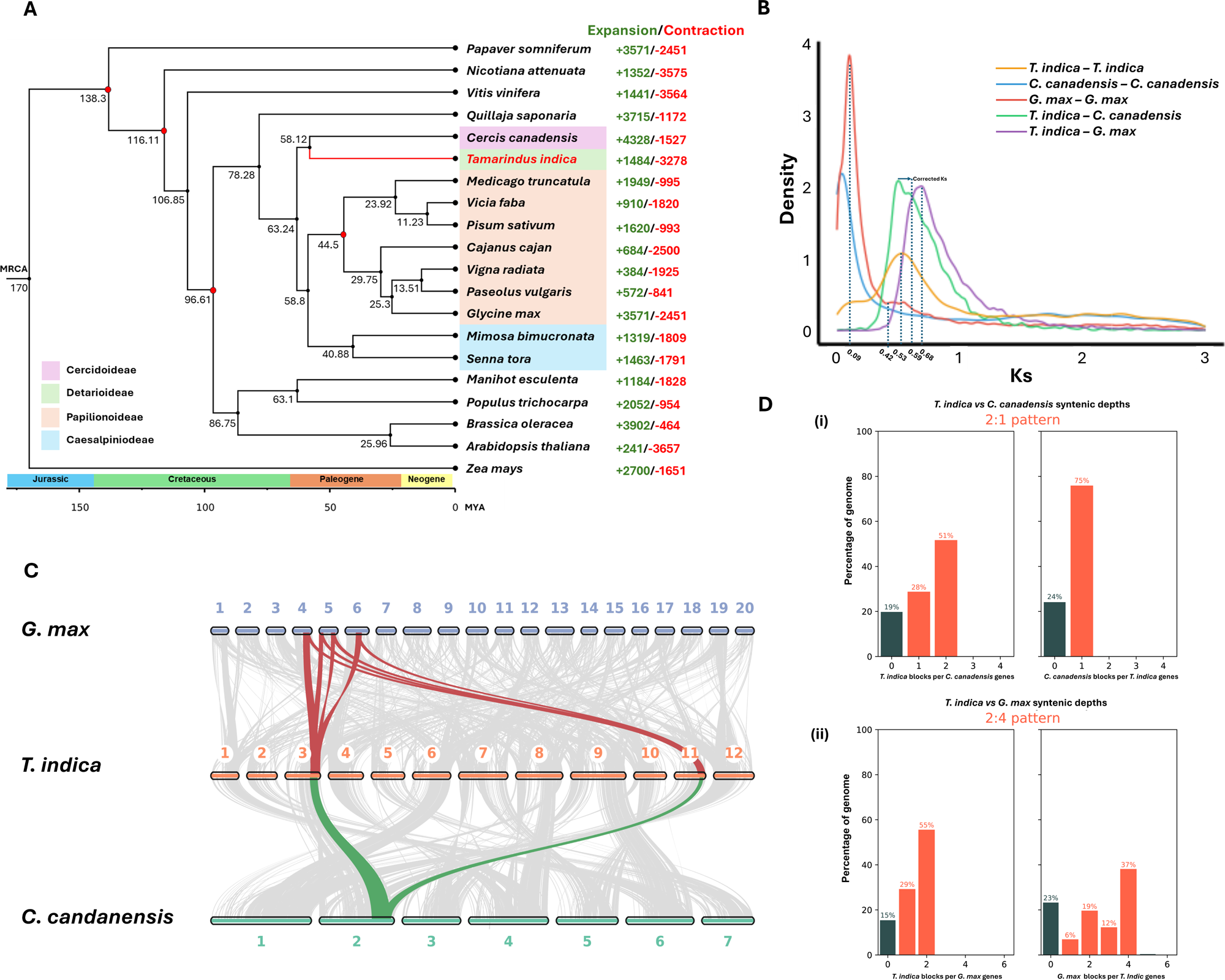
Evolutionary history of *T. indica*. **A.** Phylogenetic tree with gene family evolution. Calibrated nodes are highlighted with red dots. MRCA—most recent common ancestor, **B.** Synonymous substitution rate (Ks) distributions of paralogs and orthologs of *T. indica*, *C. canadensis* and *G. max,* **C.** Macrosynteny between *T. indica*, *C. canadensis* and *G. max*. Two gene blocks in *T. indica* exhibit a single copy in *C. canadensis* (green lines), and one gene block in *T. indica* exhibits four copies in *G. max* (red lines), suggesting that *T. indica* has undergone one independent polyploidization event after the divergence, **D.** Syntenic depth (i) *T. indica* vs *C. canadensis* (ii) *T. indica* vs *G. max*.

### Whole genome duplication (WGD) and genome synteny of *T. indica*

The Ks distribution of paralogs of *T. indica* showed a recent peak at Ks ∼0.53, which is after the divergence peak from *G. max* (Ks ∼0.68) and *C. canadensis* (Ks ∼0.59), indicating the independent whole genome duplication event in *T. indica* (**Figure 2B**). Further, the syntenic depth and whole genome dot plot analysis revealed a 2:4 syntenic ratio with *G. max* and a 2:1 ratio with *C. canadensis*, respectively (**Figure 2C, D**). These results further support that *T. indica* experienced one independent WGD, *G. max* underwent two independent WGD (Ks ∼0.42 and Ks∼0.09), and *C. canadensis* did not experience any WGD post-divergence (11,12).

Furthermore, intraspecific genome synteny of *T. indica* revealed 27% collinearity. Whereas interspecific genome synteny between *T. indica* and *G. max* was 32% and between *T. indica* and *C. canadensis* was 30%, suggesting a high collinear and syntenic gene order with other legumes (**Figure 2C**).

### Landscape of WGD/Segmental duplicates and their expression divergence in *T. indica* genome

*T. indica*, after divergence, underwent an independent WGD, which resulted in whole genome duplicates/segmental duplicates (here onwards, referred to as ‘segmental duplicates’). These segmental duplicated genes are known to play a crucial role in shaping the genome evolution (13,14). In the *T. indica* genome, these genes account for 5.15% of the genome and 21.10% of total protein-coding genes. Further, 97.5% of these segmental duplicates were present between chromosomes (Inter), which was much greater than the percentage of segmental duplicates (3.5%) occurring within the chromosomes (Intra) (**Figure 3A** and **3B**). KEGG enrichment analysis revealed that these segmentally duplicated genes were mainly involved in various metabolic pathways, including secondary metabolites biosynthesis, plant hormone signal transduction, plant-pathogen interaction, phenylpropanoid biosynthesis, etc. (**Figure 3C**).

**Figure 3:**
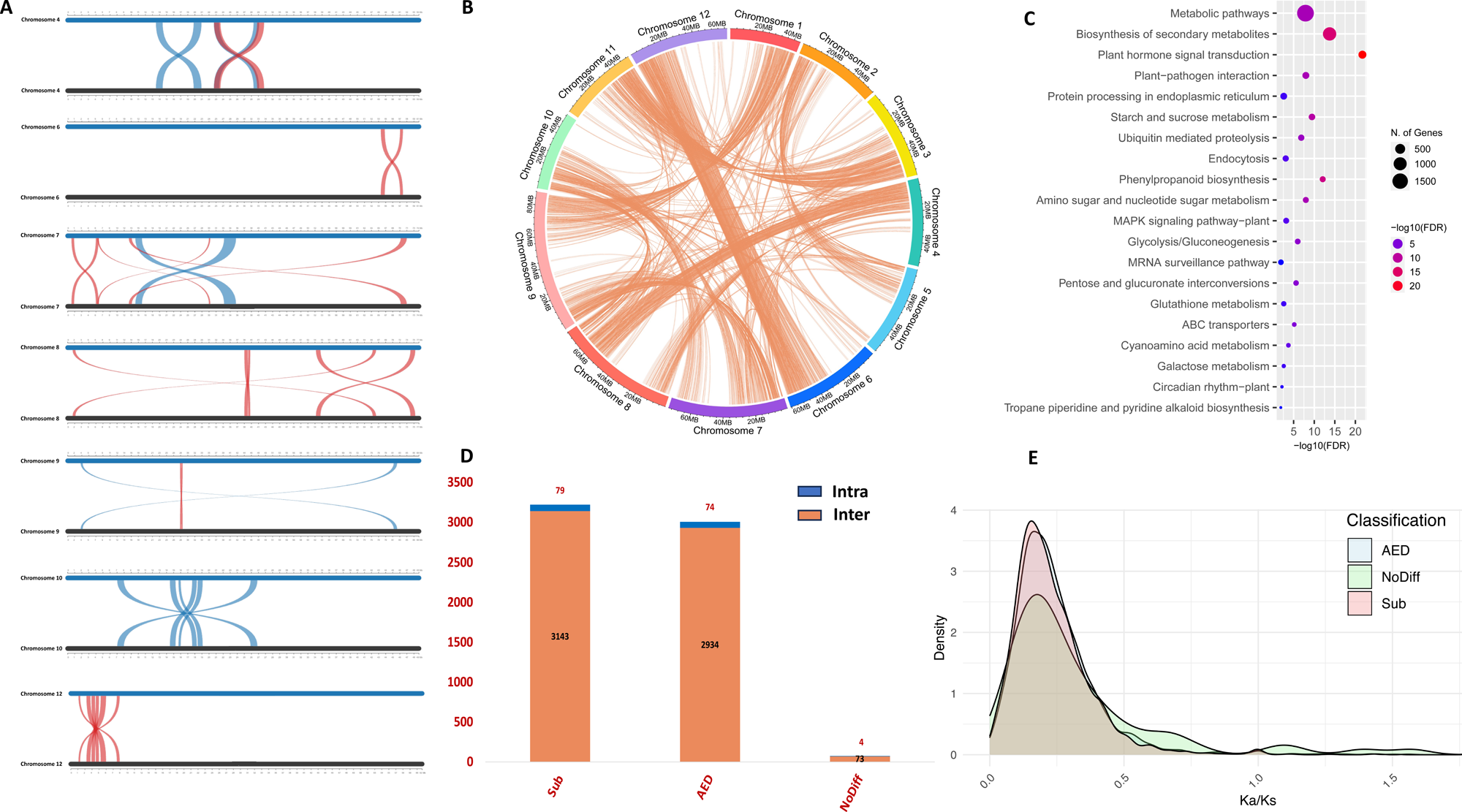
WGD/Segmental duplication and expression analysis. **A.** Distribution of intrachromosomal segmental duplication. Blue and red represent forward and reverse matches, respectively, **B.** Distribution of interchromosomal segmental duplication, **C.** KEGG enrichment of segmental duplicates (adjusted *P-*value <0.05), **D.** Stacked columns chart showing the ratio of intra to interchromosomal segmental duplicates distribution in AED, Sub, and NoDifff segmental duplicated gene pairs categories, **E.** Distribution of Ka/Ks ratio of AED, Sub, and NoDifff segmental duplicated gene pairs. Note: Gene pairs of the chromosome regions are considered.

WGD in land plants is widely accredited as a key driver of evolution and diversification. The gene duplications arising from WGD provide opportunities for sub-functionalization and neo-functionalization, enhancing adaptability and evolutionary potential (15). Based on the classification criteria (see methodology), of the 6,307 segmentally duplicated gene pairs, 3,222 were categorized as Sub, 3,008 as AED, and 77 as NoDiff class. Further, Sub, AED, and NoDiff classes have 79, 74, and 4 gene pairs found on the same chromosomes (Intra), respectively (**Figure 3D**). The Ka/Ks peak value of most of the segmentally duplicated gene pair classes was found to be less than 0.5, which suggests that a significant fraction of these gene pairs arose because of the recent WGD event of *T. indica* and are under strong purifying selection (**Figure 3E**).

## Discussion

*Tamarindus indica* is a significant horticultural crop with a wealth of pharmacologically important metabolites and nutraceuticals. However, the lack of a reference genome limits the genetic studies, the investigation of important genes and key biosynthetic pathways of this plant. Thus, in this study, we report a high-quality chromosome-scale genome assembly of *T. indica*, with an N50 of 56.6 Mb and an LAI score of 10.84, indicating the assembly to be of reference standard. The genome was found to have 1.12% heterozygosity with 63.85% repetitive sequences.

*T. indica* belongs to the third largest angiosperm family, the Leguminosae. Previously, it was placed in the Caesalpinoideae subfamily, but in 2017, the Legume Phylogeny Working Group revised the higher-level classification of the legume family and formed Cercidoideae, Detarioideae, Dialioideae, and Duparquetiodeae as distinct subfamily and placed *T. indica* into Detarioideae (10). Our phylogenetic analysis also placed *T. indica* in a separate clade from the Caesalpinoideae subfamily and closest to *Cercis canadensis* of the Cercidoideae subfamily.

Ks analysis revealed that *T. indica* experienced an independent WGD after splitting from Cercidoideae (*C. canadensis*) at Ks peak ∼0.53. Synteny analysis corroborates these findings, with a 2:1 syntenic pattern observed between *T. indica* and *C. canadensis*, indicative of a recent WGD event in *T. indica* and the absence of any independent WGD in *C. canadensis* post divergence, which is also reported in the recent genomic studies of the genus *Cercis* (11,16). The resulting gene duplicates from WGD confer genes with potential sub-functionalization and neo-functionalization, contributing to better adaptivity of the plant. The *T. indica* genome revealed 6,307 gene pairs with segmental duplications, with over half categorized as sub-/neo-functionalized pairs, while nearly the other half exhibited asymmetric expression patterns. Both categories exhibit genes involved in key pathways, such as plant hormone signaling, carbon fixation, plant-pathogen interaction, etc., mediating plant growth and resilience with biotic and abiotic stresses. Multiple duplicated copies of these functionally essential genes are potentially associated with ecological adaptations (17,18). In light of these findings, our result suggested an evolutionary fitness advantage to *T. indica*, offered by these duplicates arising from the recent WGD event.

In conclusion, our work presents the first high-quality genome assembly of *T. indica,* providing insights into its evolutionary history and genome duplication. The identification of recent WGD and the analysis of segmental duplicates highlights the adaptive potential of the plant. As *Tamarindus* is a monotypic genus, this study holds significance by offering comprehensive genomic clues that contribute to the unique traits of the plant and lay the foundation for future exploration.

## Methodologies

### DNA and RNA extraction and sequencing

Leaves of the plant were collected from the tree growing in the campus of IISER Bhopal (23.2° N 77.2° E). Species confirmation was done using the molecular barcodes ITS and matK. High molecular DNA was extracted from fresh leaves using CTAB lysis buffer, as described in Bisht et al. (13). DNA purity and concentration were checked using NanoDrop™ 8000 Spectrophotometer (ThermoFisher Scientific, USA) and Qubit 2.0 fluorometer. The 10x genomic library was prepared on the Chromium Controller instrument (10x Genomics, CA) and sequenced on the Illumina Novaseq 6000 platform to generate 60Gb data. For Nanopore sequencing, a few rounds of purification and size selection were performed using AMPure XP magnetic beads (Beckman Coulter, USA). DNA with optimum quality was taken forward for library preparation using the ligation kit SQK-LSK110 (Oxford Nanopore), and sequencing was performed on flowcell FLO-MIN106 on the MinION Mk1C platform (MinKNOW software version 22.08.6).

For RNA extraction, leaves and fruits were collected in February, a few weeks before the onset of ripening of the fruit. The leaf, fruit pulp, and seeds were immediately transferred to the laboratory, flash frozen, and stored at -80°C till further processing. The flowers were collected in April, flash-frozen, and stored at -80°C. For the RNA extraction, 1 g tissue was crushed in liquid nitrogen and transferred to 10ml prewarmed lysis buffer (300mM Tris HCl, 25mM EDTA, 2M NaCl, 2% CTAB, 2%PVP) (19). 2% beta-mercaptoethanol was added to the lysis buffer just before the lysis procedure. Tissues were lysed for 15 minutes at room temperature with intermittent mixing by inverting the tubes every 3-4 minutes. Two rounds of chloroform: isoamyl alcohol extraction was performed, followed by the addition of 0.3x volume of sodium acetate (3M, pH=5) and 0.6x volume ice-cold isopropanol. The tube was kept at -20°C for efficient precipitation, and the pellet was collected after 3 hours by centrifugation at 5000g for 30 minutes at 4°C. The pellet obtained was washed with 70% ethanol and suspended in 1 ml Nuclease-free water. 0.3x Volume of 8M LiCl was added, followed by overnight incubation at -20°C for RNA precipitation. RNA was pelleted by centrifugation at 20000g for 30 minutes at 4°C and eluted in NFW after ethanol washing. Further purification of the RNA was done using the Qiagen RNeasy mini kit spin column following the manufacturer’s protocol, with on-column DNA digestion (20). The RNA quality was assessed on NanoDrop 8000 Spectrophotometer (ThermoFisher Scientific, USA), and quantification was done on Qubit 2.0 fluorometer using Qubit RNA BR Assay kit (Invitrogen, USA). The library preparation was carried out using the QIAseq® Stranded RNA Library Kit. Final libraries were quantified, and insert size was determined using Tapestation 4150 (Agilent) utilizing sensitive D1000 screencaps (Agilent). The library with passed QC was taken forward for sequencing on NovaSeq 6000, generating 15Gb data per tissue.

### Hi-C library preparation and sequencing

The Hi-C library was prepared with young and freshly collected leaves of the plant using the EpiTect Hi-C kit (Qiagen), following the manufacturer’s protocol. The crushed leaves were fixed with formaldehyde to crosslink the chromatin inside the nucleus. Following cell lysis, intact nuclei were collected and digested using the Hi-C digestion solution provided in the kit. Sticky ends were labeled and ligated, followed by de-crosslinking and DNA purification. The extracted DNA was quantified and fragmented to 450-500 bp length. The labeled Hi-C fragments were captured via streptavidin beads pulldown. Subsequent library preparation was carried out using the standard procedure of end-repair, phosphorylation, and dA tailing of Hi-C fragments. After Illumina adapter ligation, the insert size of the prepared library was determined on Tapestation 4150 utilizing high-sensitive D1000 screentapes (Agilent). The sequencing was done on the Illumina NovaSeq 6000 platform.

### Data preprocessing and Genome-size estimation

The barcoded 10x Genomics linked reads were trimmed using the proc10xG python scripts (https://github.com/ucdavis-bioinformatics/proc10xG) to remove the barcode sequences. Nanopore reads were basecalled using Guppy v3.2.1 (Oxford Nanopore Technologies), and adapters were trimmed using Porechop v0.2.4 (Oxford Nanopore Technologies).

The 10X trimmed reads were used for genome size estimation of *T. indica* using k-mer-based analysis. The k-mer frequencies (m=21) of the raw reads were calculated using the Jellyfish v2.3.1 (21) and used for the estimation of genome characteristics using GenomeScope2.0 (22).

### Genome assembly and quality assessment

The 10x linked reads were assembled using Supernova v2.1.122 with the ‘maxreads=all’ parameter. The assembly was corrected with the help of barcode-filtered reads (https://support.10xgenomics.com/genome-exome/software/pipelines/latest/installation) using Tigmint v1.2.6(23) and scaffolded using ARCS v1.1.1(24) with default parameters. Adapters-free nanopore reads were used to further scaffold the genome assembly using LINKS v1.8.6 (25) with default parameters.

Long read assembly of the pre-processed Nanopore reads was performed using Flye v2.9.1 (26) with default parameters. The barcode-flitered10x Genomics linked reads were used for polishing (three iterations) and scaffolding the long-read assembly using Pilon v1.23 (27) and ARCS v1.1 (24), respectively. Nanopore long reads were used for further scaffolding of the assembly using LINKS v1.8.6 (25). The constructed 10x Genomics assembly and Nanopore assembly of *T. indic*a were merged using Quickmerge v0.3 (28) to achieve a more contiguous assembly. LR_Gapcloser (29) was used in five iterations for closing the gaps in the merged assembly using adapter-free long reads, and the final polishing of the assembly was performed using Pilon v1.23 (27). Purge Haplotig v1.1.270 (30) was used to remove redundant heterozygous regions responsible for increased assembly length.

To obtain the chromosome-level genome assembly of *T. indica*, clean Hi-C paired-end reads were aligned to draft assembly using BWA (31) and a deduplicated list of Hi-C reads was generated using the Juicer pipeline, and the draft assembly was scaffolded using 3D-DNA (32). The resulting assembly was then visualized and manually curated using Juicebox v1.11.08 software (33). The gaps between the pseudochromosomes were then filled by TGS-Gapcloser v1.2.1 (34) by two rounds using long-read data.

The pre-processed raw 10x linked reads and Nanopore reads were mapped to the assembled genome using BWA-MEM (v0.7.17) (35), Minimap2 (v2.17) (36), respectively, and the mapping statistics were calculated using SAMtools (v1.9) (37) “flagstat” utility, to validate the quality of the assembly. Further, the completeness of the genome assembly was analyzed through Benchmarking Universal Single-Copy Ortholog (BUSCO) v5.4.3 (38) with the embryophyta_odb10 database. Also, the LTR Assembly Index (LAI) was calculated to assess the continuity of Long Terminal Repeat retrotransposons (LTR-RTs) in the *T. indica* Genome using LTR_retriever v2.9.0 (39).

### Gene set construction and annotation

The repetitive sequences in the genome of *T. indica* were identified, and a *de novo* repeat library was constructed using RepeatModelar v2.0.3(40). The redundant sequences were clustered using CD-HIT-ESTv4.8.1 (41) with a seed size of 8bp and a sequence similarity threshold of 90%. The obtained repeats were soft masked in the genome assembly using RepeatMasker v4.1.2 (http://www.repeatmasker.org), and the masked genome assembly was then used for gene set construction of the tamarind genome using the MAKER pipeline (42). A combination of evidence-based and *ab initio* gene prediction models was used in the MAKER pipeline in three iterations. Transcriptome assembly of *T. indica* was constructed from the quality filtered RNAseq reads of different tissues (leaf, fruit pulp, seed, flower bud) using Trinity v2.14.0 (43). This *de novo* transcriptome assembly of tamarind and protein sequence of 12 Leguminosae species (*Acacia crassicarpa, Cajanus cajan, Glycine max, Lupinus angustifolius, Medicago truncatula, Mimosa bimucronata, Phaseolus vulgaris, Pisum sativum, Senna tora, Trifolium pratense, Vicia faba* and *Vigna radiata*) were used as empirical evidence for the gene set construction using BLAST (44) and Exonerate v2.2.0 (https://github.com/nathanweeks/exonerate). In the last two rounds of the MAKER pipeline, AUGUSTUS v3.2.3 (45) and SNAP v1.0 (46) were used for ab initio gene prediction. The MAKER-derived gene set obtained was further refined based on Annotation Edit Distance (AED) value and length-based criteria, keeping genes with AED value <0.45 and length >150bp to obtain the final high-confidence gene set.

### Phylogeny and gene family evolutionary analysis

For the phylogenetic tree construction and analysis of the evolutionary pattern of gene families in *T. indica*, protein sequences of 10 legume species and eight species from other representative taxa were used, with *Zea mays* as an outgroup. The longest isoform of each protein was selected. OrthoFinder v2.5.4(47) was used to construct the orthogroups, followed by KinFin v1.2 (48) to extract the fuzzy one-to-one orthogroups, which were individually aligned using MAFFT v7.467 (49). The empty sites from the multiple sequence alignment were removed using BeforPhylo v0.9.0 (https://github.com/qiyunzhu/BeforePhylo), and a maximum likelihood phylogenetic tree was constructed using RAxML v8.2.12 (50) with the ‘PROTGAMMAAUTO’ amino acid substitution model and a bootstrap value of 100. Species divergence was estimated with MCMCtree implemented in the PAML package v4.10.6 (51) using four calibration points obtained from the TimeTree database v5.0 (https://timetree.org/).

The gene family expansion/contraction analysis was performed by CAFÉ v5 (52,53). An All-versus-All mode BLASTP was performed on the longest isoforms protein sequences, and the results were clustered using MCL v14-137 (54). Filtering of gene families with genes from <2 species and gene families containing ≥100 gene copies for ≥1 species was done. The constructed species tree was converted to an ultrametric tree using r8s software (55) with a divergence estimate of 117 MYA (Million Years Ago) between *T. indica* and *V. vinifera* obtained from the TimeTree database (https://timetree.org/). Two-lambda (λ) model (birth-death rate) of CAFÉ (52) was used to analyze the evolution of gene families using the filtered gene families output and the ultrametric species tree. Gene families with ≥10 genes among the expanded gene families were considered highly expanded.

### Analysis of whole genome duplication (WGD) and genome synteny

The Ks (Substitutions per synonymous site) distribution analysis was performed to estimate WGD events in *T. indica, C. chinensis,* and *G. max* using wgd v2.23 (56). The intragenomic and intergenomic synteny was performed by MCScanX (57) with at least five gene pairs required per syntenic block, and JCVI (58) was employed to perform syntenic depth calculation, whole genome dot-plot, and macrosynteny visualization.

### Identification of WGD/Segmental duplication and expression analysis

The WGD/segmental duplicate genes were identified using the duplicate_gene_classifier implemented in MCSCanx (57). Further, collinear gene pairs present in the chromosome regions were obtained from the MCScanx result performed during synteny analysis. The TPM (transcripts per million) values of the segmentally duplicated gene pairs were calculated using Kallisto v0.48.0 (59) for each tissue. The gene pairs based on their expression (TPM value) were then classified into three categories: (i) sub-/neo-functionalized pairs (Sub), where each duplicate showed higher expression than the other in at least one sample; (ii) asymmetrically expressed duplicates (AED), where one duplicate exhibited higher expression in at least one-third of the samples and was never expressed lower than its partner in any sample; and (iii) the remaining pairs, categorized as no-difference (NoDiff) pairs (14,60). The numbers of non-synonymous substitutions per synonymous site (Ka), synonymous substitutions per synonymous site (Ks), and the Ka/Ks ratios were calculated using KaKs Calculator 3.0 (61).

## Acknowledgments

MS and SM thank Council of Scientific and Industrial Research (CSIR) for the fellowship. MSB thanks Ministry of Education, Govt. of India for Prime Minister Research Fellowship (PMRF). The authors also thank the Sequencing and MASS facility at IISER Bhopal, and thank the Department of Biotechnology for the financial support (BT/PR51934/BTIS/137/86/2024) and the intramural research funds provided by IISER Bhopal.

## Authors’ contributions

VKS conceived and coordinated the project. MS performed sample collection, DNA-RNA extraction, prepared samples for Hi-C sequencing, and performed Nanopore sequencing. SM performed DNA extraction and sample preparation for 10x sequencing. MSB performed the computational analysis of the study. MS and MSB analyzed the data and interpreted the results. MS and AMG performed metabolome analysis. MS and MSB constructed all the figures. MS, MSB, and VKS wrote the manuscript. All the authors have read and approved the final version of the manuscript.

